## Supporting Information for "Potential Neutralizing Antibodies Discovered for Novel Corona Virus Using Machine Learning"

**Supplementary Figure 1.** Contact Distance plot of trajectories of all the 22 mutant structures and the crystal structures 2GHW and 6NB6. The contact distances were computed using MDTraj package. Instantaneous contact distances for each timestep are shown for the course of the simulation (15 ns). The contact distance values vary within a small range for all the structures, indicating that simulations did not have any abrupt changes.

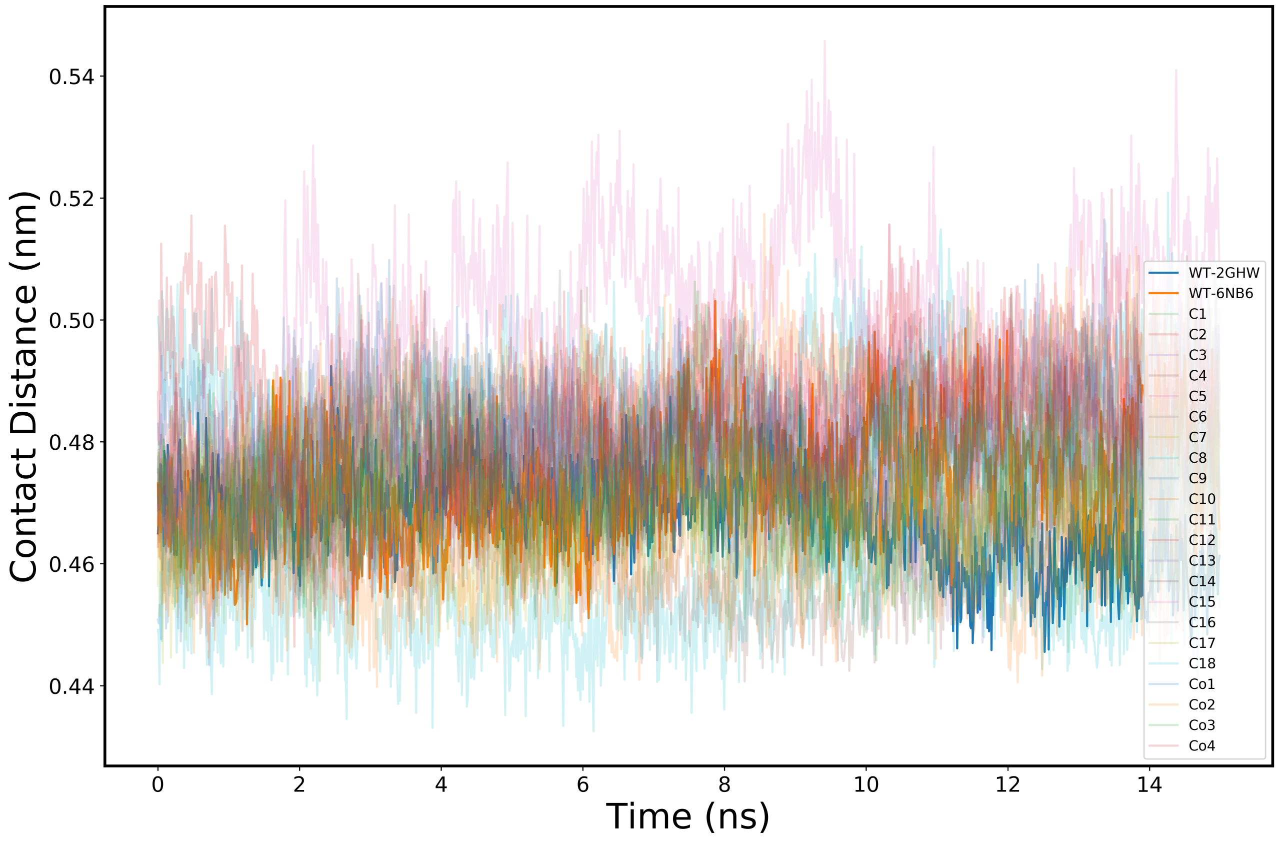

**Supplementary Figure 2.** Root Mean Square Deviation (RMSD) plot of trajectories of all the 22 structures and the crystal structures 2GHW and 6NB6. The deviations are within acceptable range indicating the strucutres were stable over the course of simulation. RMSD was averaged over the entire simultation and subsequently used for generation of Figure 4b.

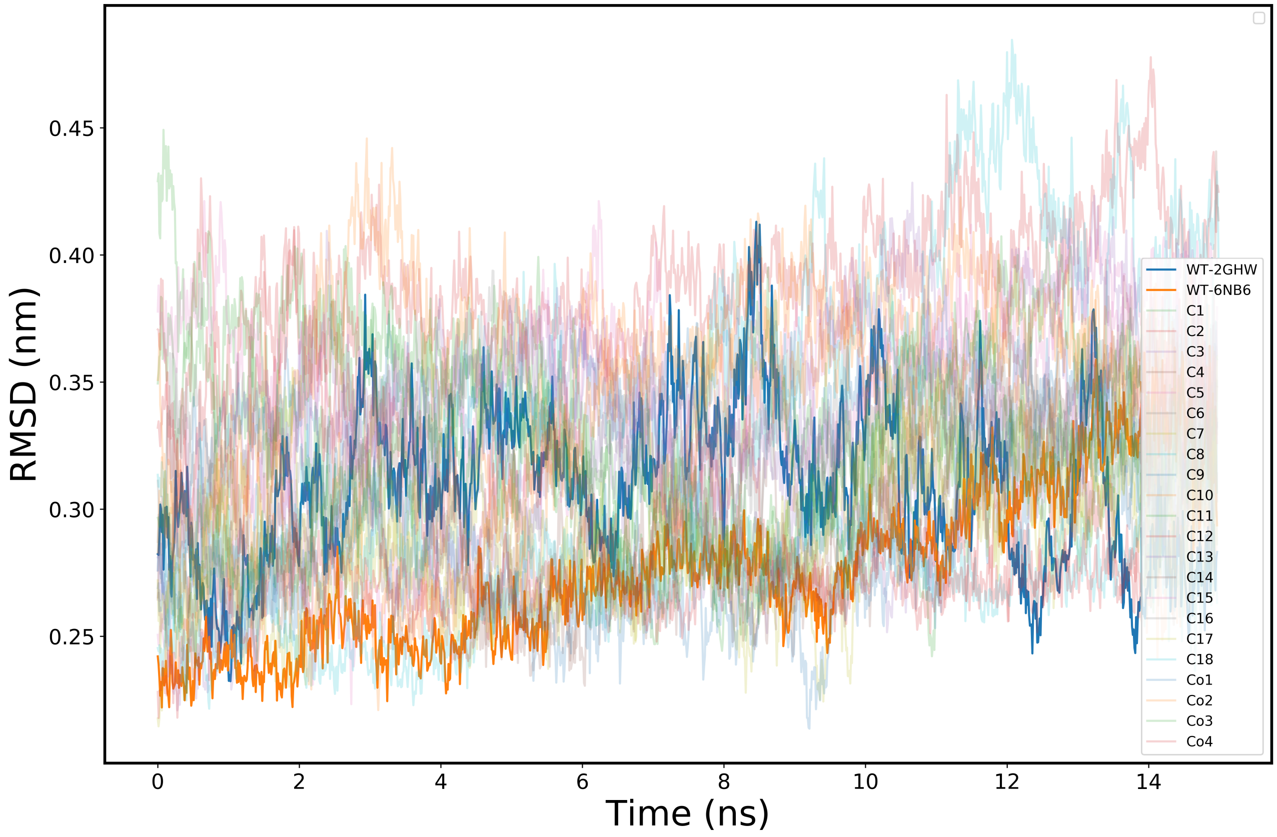

**Supplementary Figure 3.** Structure of virus antigen and antibody along with the epitope interactions in the crystal structure PDB ID: 2GHW. The viral antigen is in green and antibody is in cyan. Black lines indicate electrostatic interactions between the antigen epitope and the antibody. These interactions play a significant role in the specificity of antigen recognition by antibody.

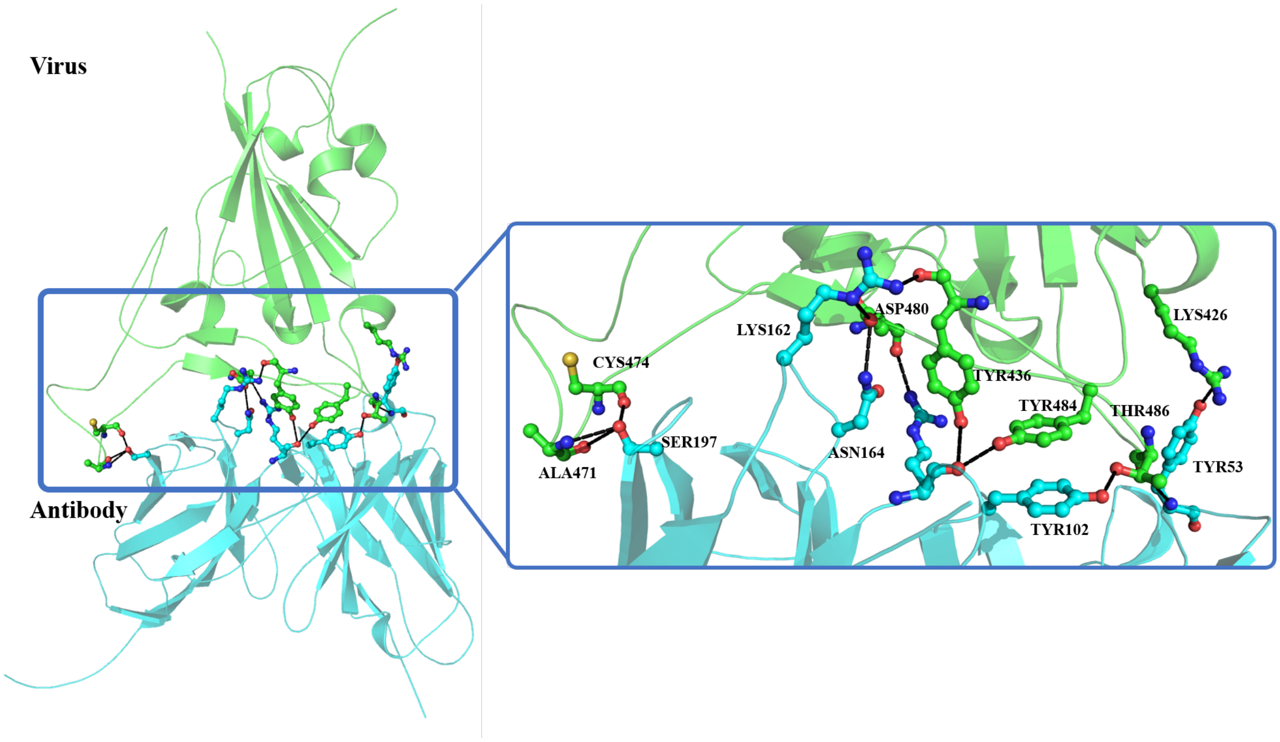

**Supplementary Figure 4.** Native contacts between the antibody and antigen in the crystal structure (PDB ID: 2GHW) shown pictographically. The amino acids above black dotted line represents the antibody (pink) and below it is the viral epitope (brown). Electrostatic interactions are represented by the whole amino acid ball-stick diagram and are connected by green dotted lines.

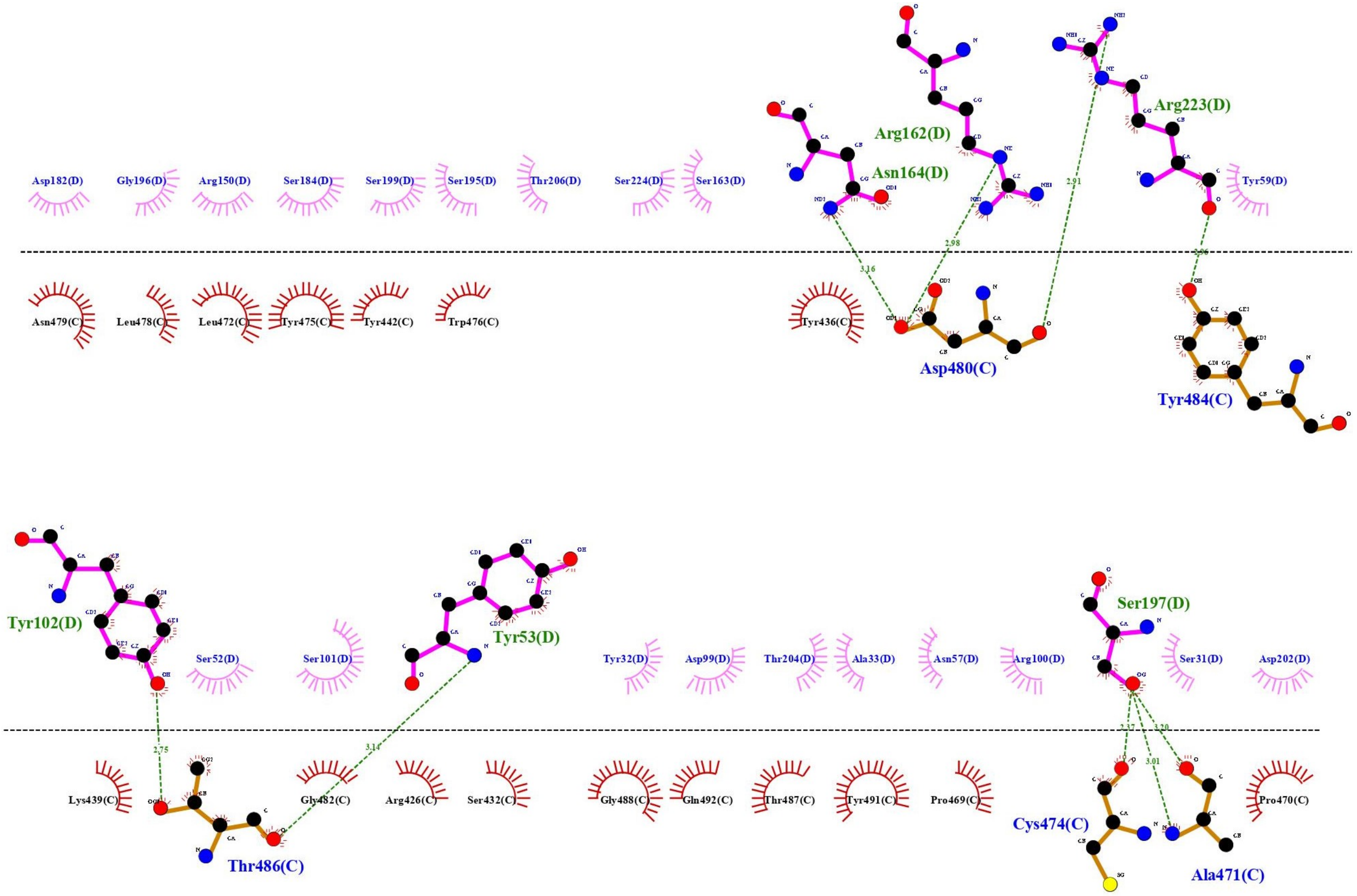

**Supplementary Figure 5.** Predicted contacts between the antibody and COVID-19 antigen. The COVID antigen is overlaid onto the crystal structure antigen (2GHW) to conserve the geometry and make predictions. The amino acids above black dotted line represents the antibody (pink) from PDB: 2GHW, and below is the viral epitope(brown) from COVID-19 sequence. Electrostatic interactions are represented by the whole amino acid ball-stick diagram and are connected by green dotted lines.

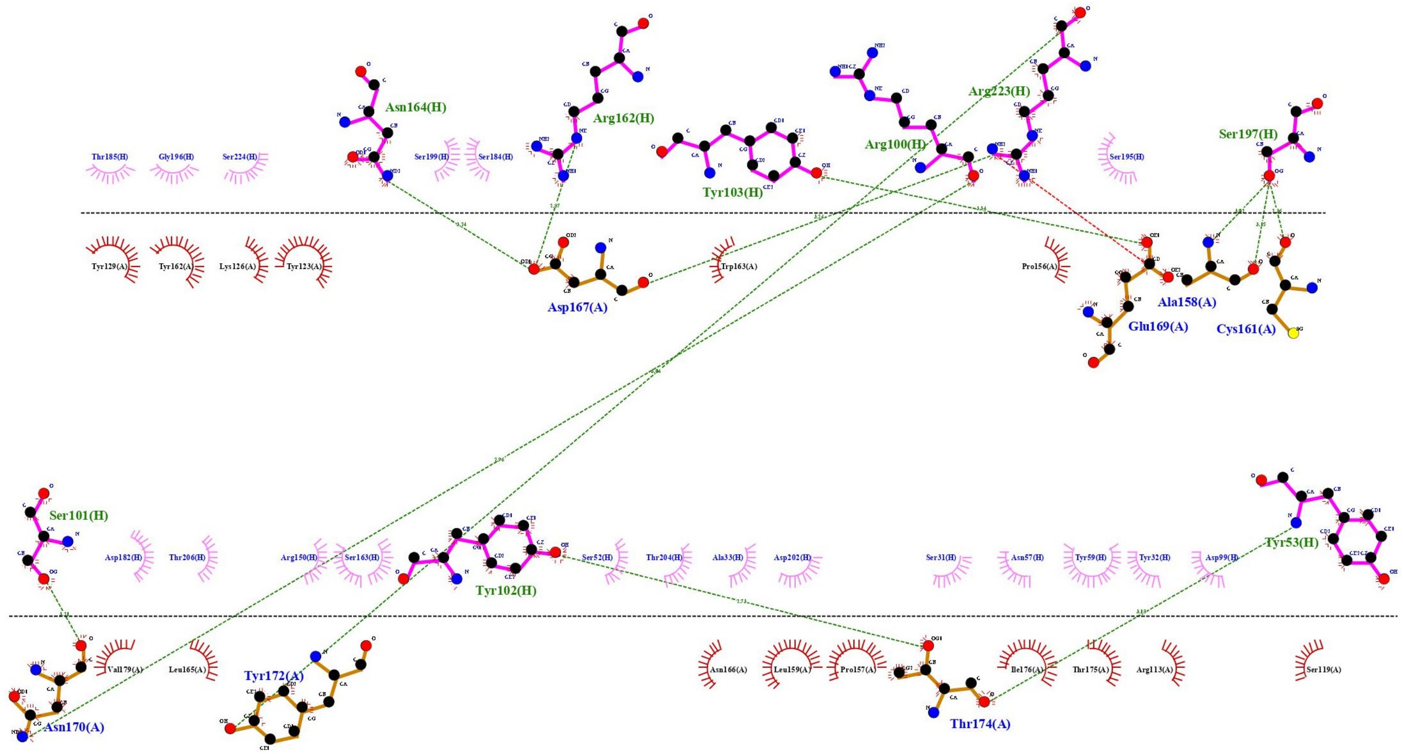

**Table S1:** The neutralizing candidates obtained before screening for stability, and after screening with ML and Bioinformatics. There are 18-point mutations in total which are predicted to have a neutralizing effect on COVID-19.

| **Structure** | **Mutation** |
|---|---|
| C1 | 2GHW – A33C |
| C2 | 2GHW – V50M |
| C3 | 2GHW – I51M |
| C4 | 2GHW – I51V |
| C5 | 2GHW – N57H |
| C6 | 2GHW – R100H |
| C7 | 2GHW – R150H |
| C8 | 2GHW – V161M |
| C9 | 2GHW – R162H |
| C10 | 2GHW – N164H |
| C11 | 2GHW – T185N |
| C12 | 2GHW – R186H |
| C13 | 2GHW – F203M |
| C14 | 2GHW – T204N |
| C15 | 2GHW – T206N |
| C16 | 2GHW – R223H |
| C17 | 6NB6 – R56H |
| C18 | 6NB6 – K58S |

**Table S2:** The possible co-mutations from 5 stable neutralizing candidates. Structures Co1 – Co4 were predicted as neutralizing, however; structure Co5 was predicted as non-neutralizing by the XGBoost model. All these co-mutant structures we MD validated for their stability.

| Structure | Mutation | Neutralization Potential |
|---|---|---|
| Co1 | 2GHW – I51M, R150H, T204N | 1 |
| Co2 | 2GHW – I51M, R150H | 1 |
| Co3 | 2GHW – I51M, T204N | 1 |
| Co4 | 6NB6 – R56H, K58S | 1 |
| Co5 | 2GHW – R150H, T204N | 0 |

**Supplementary Data.** Stable_str.zip. The zip file contains the PDB coordinates for the final frame of the simulation of each of the neutralizing candidate antibody. All the mutant antibodies are structurally intact and are stable.

### Supplementary Methods

#### $IC_{50}$ Value interpretation

The $IC_{50}$ values in dataset varied from 0 to >50. For simplicity of analysis, we classified the $IC_{50}$ values into two categories. Where, $IC_{50} > 10$ would indicate a non-neutralizing antibody and $IC_{50} < 10$ would indicate a neutralizing antibody. Many viruses had the same viral epitope and yet had very different $IC_{50}$ values. As a result, the model had to predict two different labels for the same antibody-antigen combinations. We resolved this, by taking a majority vote among the conflicts and transformed the labels which do not concur with the majority. Employing this scheme improved the results of our model.
